# An attempt to find the correlations between body weight and the composition of gut microbiota in Zhejiang and Shanghai

**DOI:** 10.1101/2022.05.05.489247

**Authors:** Yihan Xia, Ziying Jin

## Abstract

Previous studies showed that the human gut microbiota was associated with metabolic diseases, but the interaction and mechanism between the gut microbiota and metabolic disease are still unclear. In this study, the gut microbiota of 58 persons living in Zhejiang and Shanghai area will be analyzed. Then, the potential contribution of the human gut microbiota to obesity/high Body Mass Index (BMI) will be explored. The gut microbiota was studied by high throughput sequencing analysis of bacterial 16S rRNA gene fragments, and the gut microbiota samples with different BMI were compared. Meanwhile, some gut microorganisms from faeces of a healthy individual were cultivated and isolated, and the classification was identified by 16S rRNA sequencing. The main microbes in human gut microbiota were assigned to the phyla of Firmicutes, Bacteroidetes, Proteobacteria, and Actinobacteria. Moreover, four strains were isolated from an individual fecal sample, of which one species was assigned to *Escherichia fergusonii* and the other three strains were assigned to *Weissella cibaria*. These four species belong to both abundant and low-abundant species revealed by high throughput sequencing. It was found that individuals with different BMI have different gut microbiota; while the differences are not significant. Also, the Firmicutes/Bacteroidetes ratio increases with the decrease of BMI, which is corresponding to previous results. In the future, more cohort gut microbiota in Zhejiang and Shanghai area will be collected and recovered, and the gut microbiota database of Zhejiang and Shanghai area will be built up in order to provide the basis for future gut microbiota modulation in this area.

## 1. Introduction

There are a huge number of microorganisms in the human digestive system, known as human gut microbiota. The total amount of microbes in gut microbiota reaches 10^13^, which is comparable to the number of human somatic cells. Recently, numerous studies have shown that the interaction between the gut microbiota and human body would either directly or indirectly affect human health. The gut microbiota can synthesize essential vitamins and amino acids [1], help digest complex carbohydrates [2] and secrete short-chain fatty acids (SCFAs) to improve intestinal health. Probiotics in the gut suppress the growth of pathogens through colonization and secretion of bacteriocins [3]. Studies on the intestinal-brain axis have confirmed that there is two-way interaction between gut microbiota and the brain, which has an important impact on the occurrence and development of depression, etc [4].

The disorder of gut microbiota is closely related to digestive, metabolic and neurological diseases, such as inflammatory bowel disease, type II diabetes, and obesity [5]. With the deepening of the research on the correlation and causality between gut microbiota and various diseases, regulation of gut microbiota has become a choice for the treatment of various intestinal metabolic diseases. At present, methods such as fecal bacteria transplantation which transplant the gut microbiota of healthy people into that of patients, and supplement of probiotics, prebiotics, and symbiosis can reconstruct the gut microbiota of patients and are applied in the treatment of digestive system or metabolic diseases such as Crohn’s disease.

Obesity is a serious public health problem affecting about 1.9 billion people around the world [6]. It is closely related to high sugar and fat diet, living habits, stress level and genetic factors. Obesity itself will lead to a variety of health problems, such as cardiovascular disease, metabolic disease, and cancer [7]. Previous studies have shown that obesity is associated with the composition of gut microbiota, for example, the proportion of certain microbe species in gut microbiota of obese people is different from that in people with normal weight [5]. Transplanting the gut microbiota of obese patients into mice will also result in obesity [8]. Further studies have shown that short-chain fatty acids and short-chain fatty acid-producing microbes are closely related to obesity [9], and probiotics regulation is expected to prevent and reduce the incidence of obesity. However, the composition of gut microbiota in different populations is affected by a variety of factors such as diet, lifestyle, genetics, and environment, especially the gut microbiota of populations in different regions are different [23]. Therefore, on the basis of existing studies, in-depth analysis of gut microbiota in specific areas is conducive to the analysis and precise regulation of gut microbiota and provide a scientific basis for the prevention and treatment of obesity.

In this study, fecal bacteria samples from 58 individuals of different ages and weights in Zhejiang and Shanghai were collected, and high-throughput sequencing was applied to reveal the gut microbiota and characteristics of samples from different populations. At the same time, intestinal bacteria samples of healthy human feces were separated and cultured, providing a bacterial base for the precise regulation of gut microbiota by fecal bacteria transplantation and other strategies in the future.

## 2. Methods

### 2.1. Collection of different fecal bacteria samples

The research objects are mainly families living in Zhejiang province and Shanghai city, and all the selected are healthy individuals, divided into three groups. The three groups were respectively underweight group (TH group, BMI≤18.5), normal group (MI group, 18.5 < BMI≤24), and overweight group (FA group, BMI > 24). A total of 58 samples were collected. More than 10 healthy people participated in each group.

All volunteers signed the informed consent form and 0.1-0.5 g of fresh stool were collected into stool preservation solution. The samples were numbered and sent to Shanghai Ling En Company for sequencing of V3-V4 fragment of 16S rRNA gene. The primers were 341F: ‘CCTACGGGNGGCWGCAG’ and 806R: ‘GGACTACHVGGGTATCTAAT’. The sequencing results have been uploaded to the SRA database, and the SRA serial numbers of 58 samples are: SRR15851344-SRR15851361, SRR15851693-SRR15851712, SAMN21396792-SAMN21396811.

### 2.2. Analysis of the results from V3-V4 fragment of 16S rRNA gene sequencing

The dix-SEQ workflow developed by Shanghai Luojie Technology was used for data processing [26]. After obtaining the original sequencing data, the low-quality data was filtered, primer sequences were removed, and both ends were spliced to obtain effective splicing data. Then, the Usearch software was used to classify the sequences with 97% similarity for Operational Taxanomic units (OTU). On this basis, the related OTUs are classified by species, and the abundance of each OTU were analyzed. Finally, the alpha and beta diversity of each community was analyzed to obtain community information within and among samples, and the differences within relevant groups and between species were analyzed. The enterotype analysis website (https://enterotypes.org/) was also utilized to analyze the enterotype of the samples collected.

### 2.3. Collection of fresh fecal bacteria sample and separation of microbes

A fresh feces sample of about 0.5 g from a male (16 years old) was taken into a sterile EP tube, centrifuged at 10000 g for 1 min, and the supernatant was discarded. 1 mL of PBS buffer was added to the precipitate, then centrifuged again at 10000 g for 1 min, and the supernatant was discarded. 1mL PBS buffer was added into the EP tube again for re-suspension.

### 2.4. Separation and cultivation of fecal bacteria

Microbial samples in the inoculation buffer were respectively streaked or coated on LB solid medium and MRS solid medium then aerobically cultured at 37 °C or 30 °C. At the same time, in the anaerobic incubator, fecal bacteria were streaked or coated on MRS solid medium and cultured at 30 °C. After overnight culture, the colonies growing on each solid plate were selected and transformed onto fresh LB or MRS plates and continued to be cultured until monoclonal bacteria grew. After the above steps were repeated once more, monoclones (i.e. isolated pure cultures) were selected from each plate and added to LB or MRS liquid medium for culture.

### 2.5. Extraction of monoclonal DNA

1000 µL bacterial solution was added into 2mL centrifuge tube. After centrifugation, each bacterial liquid sample was extracted according to the experimental protocol of bacterial genome DNA Extraction Kit (Tiagen Biochemical Technology, Beijing, China). Especially, lysozyme was added into bacteria cultured in MRS medium to destroy the cell wall during the extraction process. After extraction, DNA extraction results were detected by electrophoresis and quantified.

### 2.6. PCR amplification and detection of 16S rRNA gene

Primers 27F (5 ‘-AGA GTT TGA TCC TGG CTC AG-3’) and 1492R (5 ‘TAC GGY TAC CTT GTT ACG ACT T 3’) and Tsingke Gold Mix (green) (Tsingke Biotech, Shanghai China) were used to amplify 16S rRNA gene in each isolated pure culture. The amplification system was 50µL, containing 1* Gold Mix mixture 45µL; primers 27F/1492R each 2 µL (10µM); DNA solution 1 µL (about 20 ng DNA). PCR amplification program was set up as: 98°C, 10s, 55°C, 10s, 72°C, 30s, 30 cycles; 72 °C, 5 min. After the PCR amplification, products were verified by 1% agarose gel electrophoresis and sent to Tsingke Biotech for sequencing. Four full-length 16S rRNA genes were obtained, with serial numbers of MZ882147-MZ882150 in the Genbank database.

## 3. Results and discussion

### 3.1. Results of sample collection

A total of 58 samples were collected from 21 groups of families, among whom 16 were children (0-16 years old), 11 were young (17-40 years old), and 30 were middle-aged (over 40 years old). There were 28 males and 30 females. The above samples were divided into 3 groups according to BMI, including 13 samples in the underweight TH group (BMI≤18.5), 27 samples in the normal MI group (18.5 < BMI≤24) and 18 samples in the overweight FA group (BMI > 24). Among the 58 people, the youngest was 2 years old, the oldest was 51 years old, and the average age was 33.00. The average ages of three groups were 19.46, 35.85 and 38.56 for TH, MI, and FA group respectively. In terms of age distribution, young people have lower BMI and middle-aged people have higher BMI, which may be related to the aging and slowing down of metabolism.

### 3.2. Alpha diversity analysis of human gut microbiota from Zhejiang province and Shanghai city

After sequencing of 58 samples, a total of 2,846,080 sequences were obtained, the sequence number of each sample was ranging from 33,092 to 67,800, and the average sequences per sample was 49,070. After clustering 58 samples at a similarity level of 97%, a total of 1545 OTUs were obtained. The number of OTUs shared by all samples was 8 (0.5% of all OTUs), and the number of OTUs that existed in 90% of samples was 39 (2.5% of all OTUs). Among 58 samples, the number of OTUs of each sample ranged from 208 to 578; The average number of OTUs in the sample was 374.

Alpha diversity of the three groups of samples were analyzed, of which the differences between OTU diversities were not significant, but the richness of all the three groups was less than Chao1. Therefore, to understand the microorganisms with low abundance value in the community, the sequencing amount together with the detection rate of low abundance out should be increased. In addition, the high dominance values for each sample indicated the presence of dominant microorganisms in these gut microbiotas. Thus, further studies of these dominant microorganisms are valuable. Simpson and Shannon indexes of the three samples were similar, which further indicated that there was little difference in gut microbiota among the three groups (Table 1).

**Table 1.**
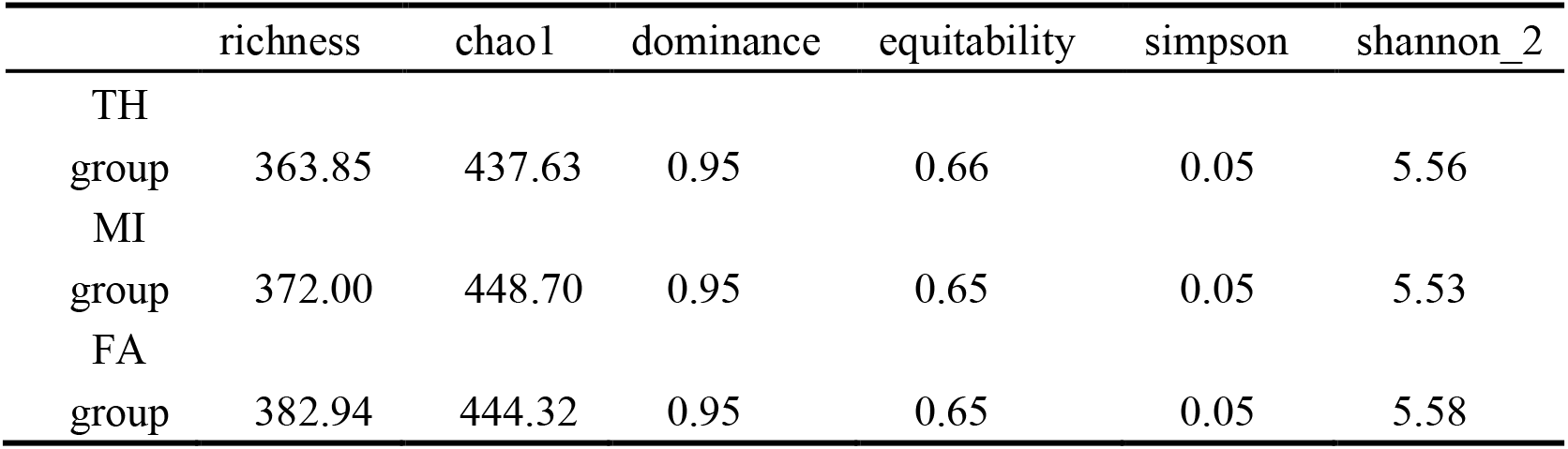
Alpha diversity of three groups of samples

### 3.3. Classification of gut microbiota at phylum level in Zhejiang and Shanghai

Firstly, bacterial community structure between different groups at the phylum and genus levels were compared. The results of phylum level showed that the intestinal bacteria in the three groups of human faeces samples were mainly composed of Firmicutes, Bacteroidetes, Proteobacteria and Actinobacteria, which together account for more than 90% of the human gut microbiota (Figure 2 and Figure 3). Among the underweight TH group, Firmicutes (36.0%) and Bacteroidetes (39.5%) were the two main types of microorganisms. There were differences in the abundance of the two types of microorganisms in the normal MI and overweight FA groups. Firmicutes and Bacteroidetes in the normal MI group accounted for 30.0% and 46.3% of the gut microbiota, respectively. In the overweight FA group, the proportion of Firmicutes and Bacteroidetes in gut microbiota was 33.2% and 47.1%, respectively (Figure 2 and Figure 3). Previous studies have shown that the ratio of Firmicutes/Bacteroidetes will decrease with the increase of BMI [16]. In this research, the results showed the ratio of Firmicutes/Bacteroidetes in the underweight group was significantly higher than those in the MI group and the FA group. There is also a certain relationship between gut microbiota and age. Firmicutes/Bacteroidetes ratio will increase first and then decrease with age [17]. As the samples were adolescent and middle-aged, Firmicutes/Bacteroidetes ratio in TH group was relatively higher; while in MI and FA groups, the ratio was lower.

**Figure 1.**
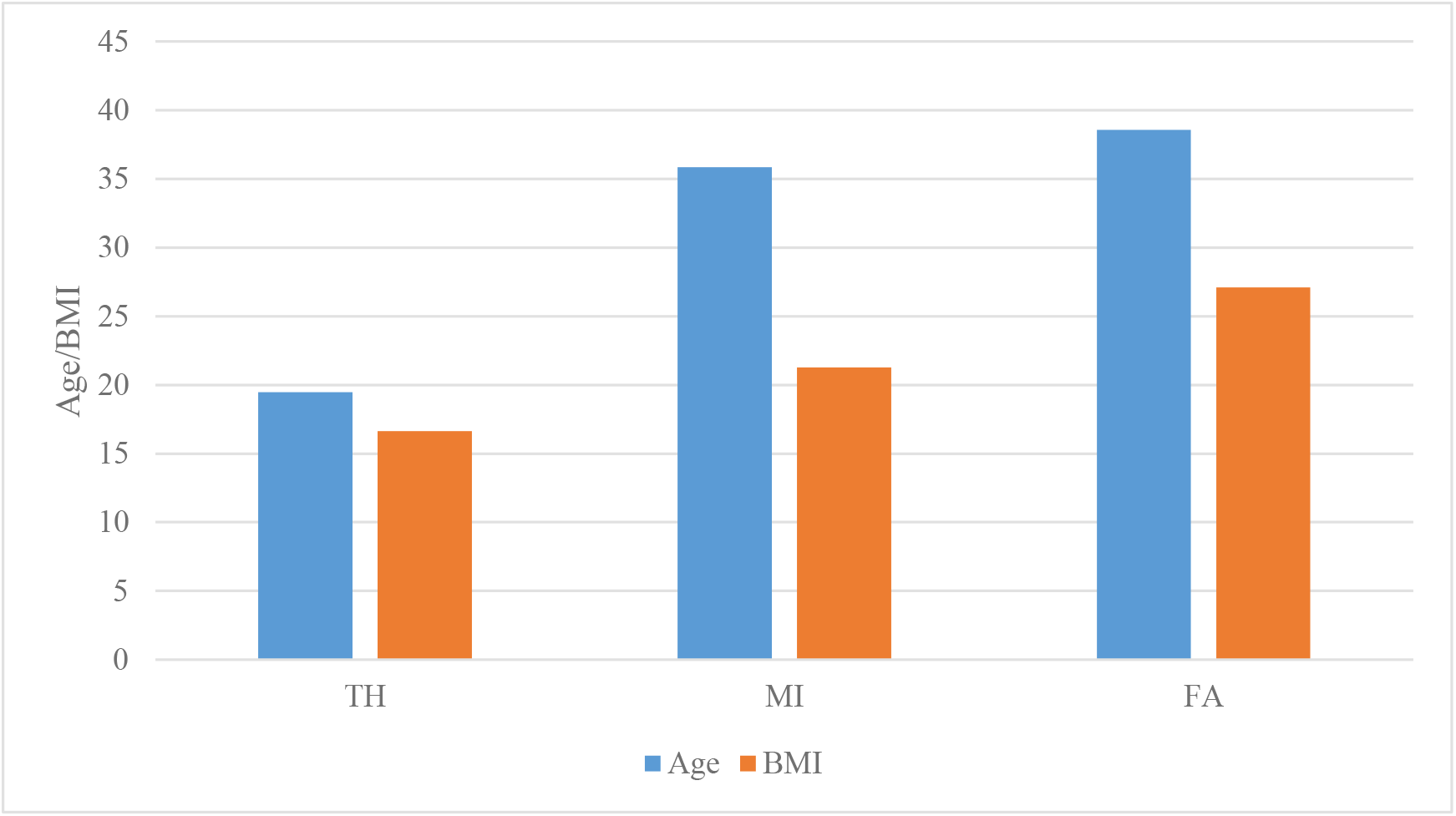
The average age and BMI of three groups

**Figure 2.**
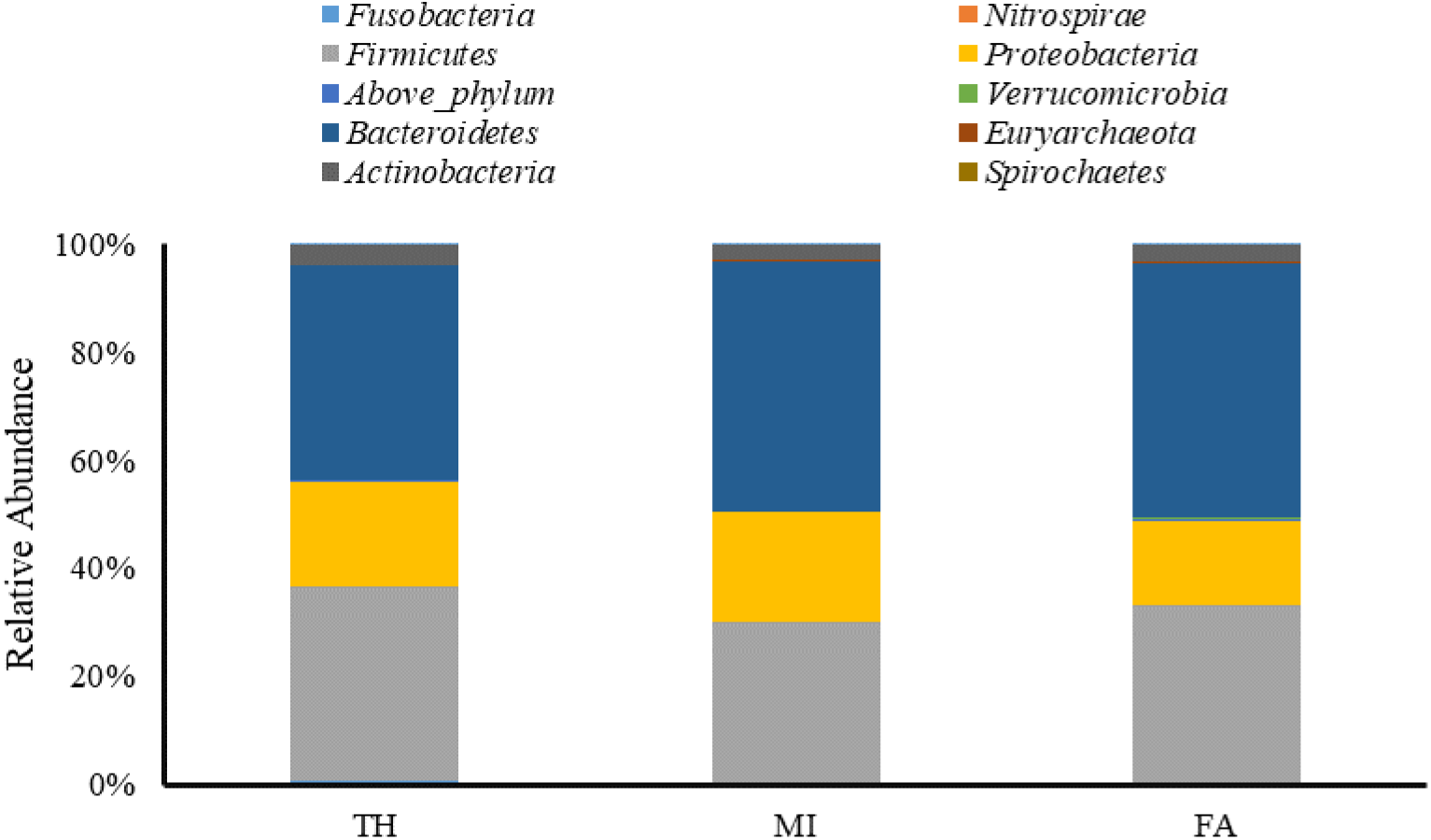
Composition of human gut microbiota at phylum level in TH, MI and FA groups in Zhejiang and Shanghai

**Figure 3.**
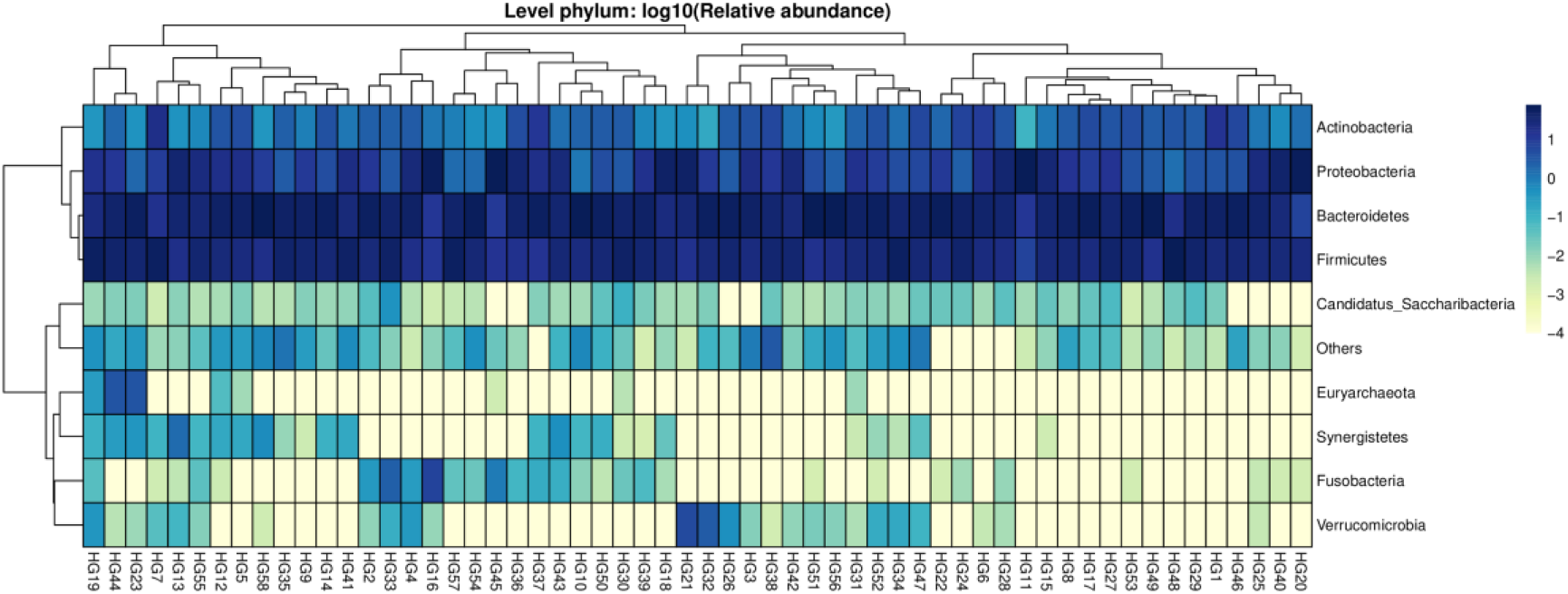
Composition of human gut microbiota at phylum level in Zhejiang and Shanghai

According to the different composition of gut microbiota, human intestinal bacteria can be divided into different enterotypes, which are related to diet and disease. After analyzing the 58 samples collected in this study, results showed there were 3 people with ET_F type, 0 people with ET_P type, 15 people with ET_B type, and 40 people with other unknown types (Figure 4). Individuals with ET_P type may have a diet mainly rich in carbohydrates such as vegetables. Individuals with ET_B type may consume more meat products which are rich in protein and fat. Individuals with ET_F type may have a diet high in dietary fiber, such as grains and fruits. Since the comparison website used in this study mainly collects gut microbiota samples from Western populations which may be less updated, hence there were many unknown enterotypes in this study. At the same time, higher BMI values of people with ET_B type were detected, further indicating that the ET_B type may be associated with higher BMI value/obesity. In the experiment, 70% of the samples had different enterotypes with existing classifications, suggesting that strengthening the establishment of the gut microbiota database in different regions of China, and improve the level of gut microbiota research in China is significant. The analysis and regulation of gut microbiota will provide evidence for future treatments of related intestinal diseases.

**Figure 4.**
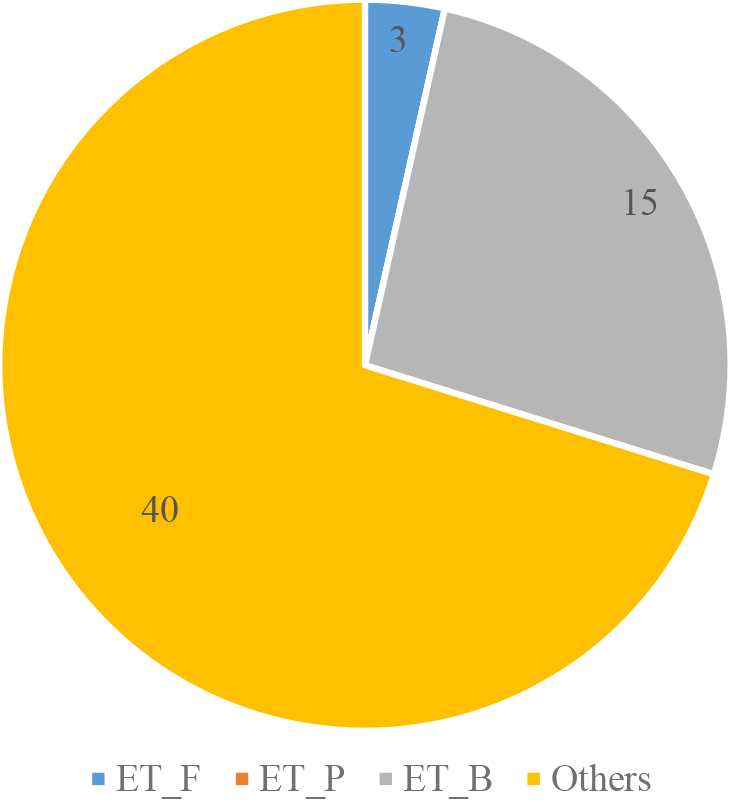
Enterotype distribution in Zhejiang and Shanghai

### 3.4. Classification of gut microbiota at genus level in Zhejiang and Shanghai

At the genus level, the dominant genera in these samples are *Bacteroides, Prevotella, Faecalibacterium* and *Escherichia* (Figure 5 and 6). *Bacteroides* is a common genus of intestinal bacteria, which can regulate human immune response, help digest and degrade polysaccharides and produce polysaccharides beneficial to human body. It can produce fatty acids as an energy source for the host through fermentation of carbohydrates, accounting for a high proportion in the industrialized population [18]. The average proportion of *Bacteroides* in 58 samples was 29.25%, and the proportion of TH, MI and FA groups was 29.00%, 30.58% and 27.44%, respectively. The average proportion of *Escherichia* in the three groups was 6.11%, and the abundance of *Escherichia* decreased with the increase of BMI in the three groups. *Prevotella* is also one of the dominant bacteria in the intestinal tract of some people in Zhejiang and Shanghai, with an average abundance of 5.49%. Its proportions in the three groups of samples were 3.57%, 4.48% and 8.38%, respectively (Figure 5). *Prevotella* have been shown to have the ability to decompose phytoglycans, and is more common in the intestinal tract of vegetarian populations [19]. In addition, the abundance of Prevotella increased with the increase of BMI, indicating that *Prevotella* may be associated with the increase of BMI (obesity), and further attempts to isolate *Prevotella* from samples and verify in vitro are of great significance .

**Figure 5.**
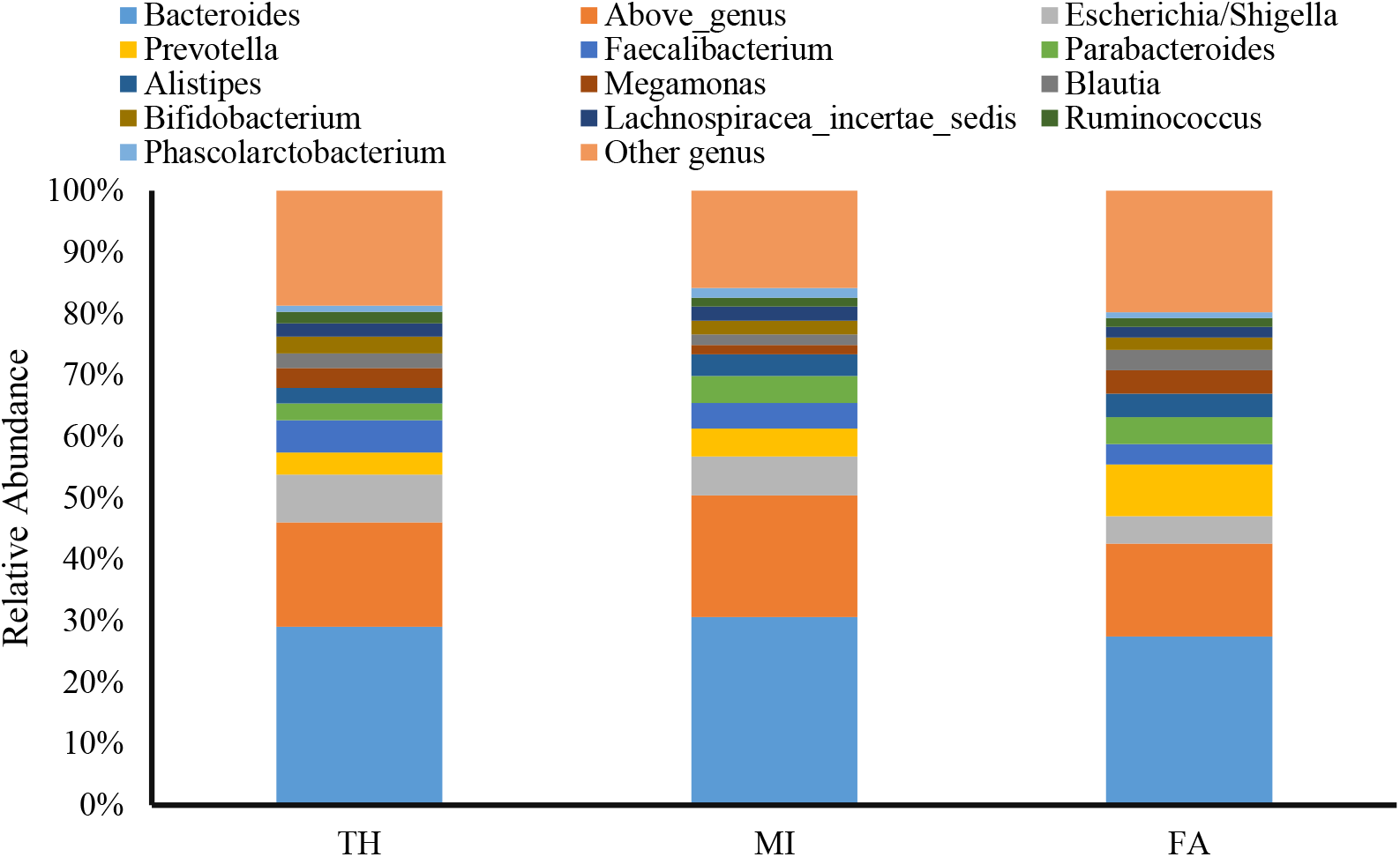
Composition of gut microbiota in different groups of people from Zhejiang and Shanghai at genus level

**Figure 6.**
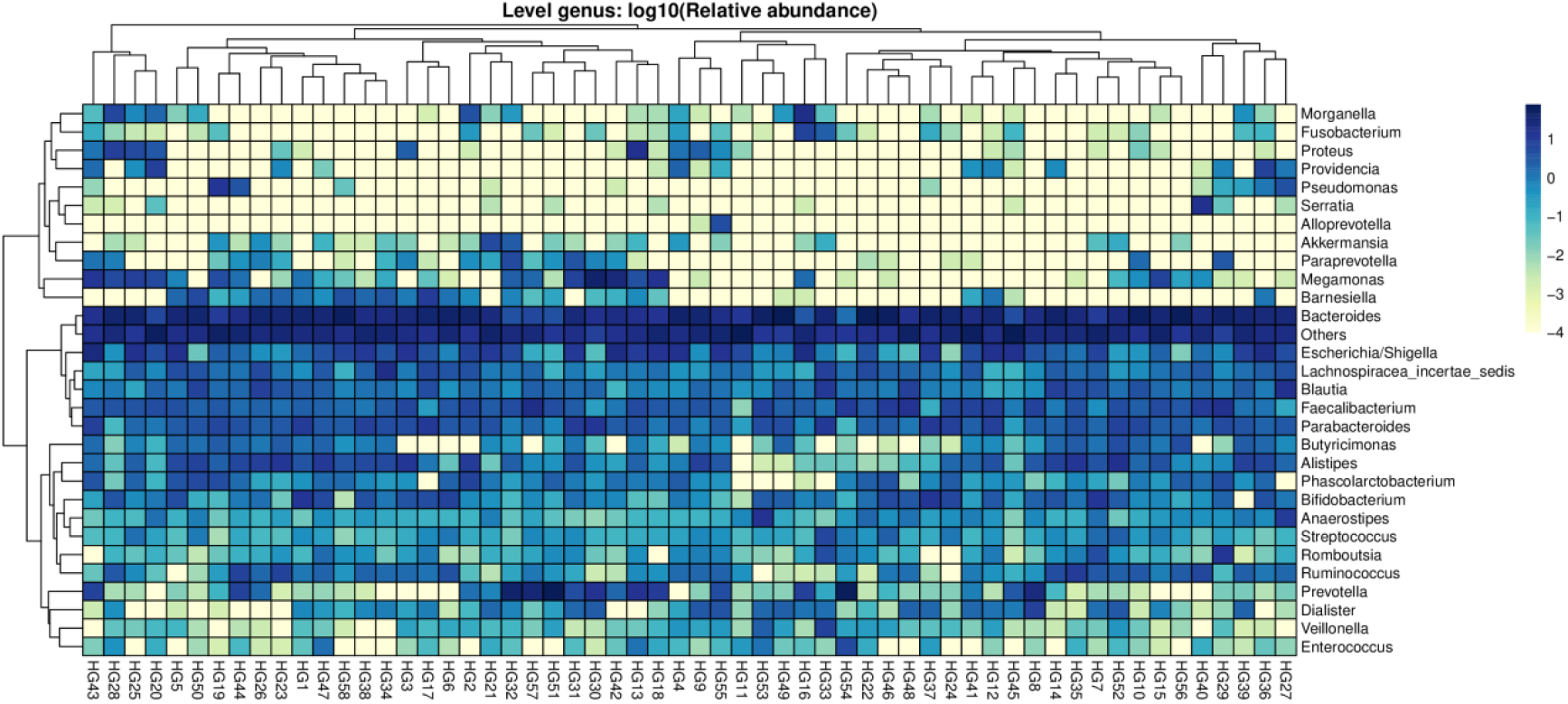
Composition of human gut microbiota at genus level in Zhejiang and Shanghai.

**Figure 7.**
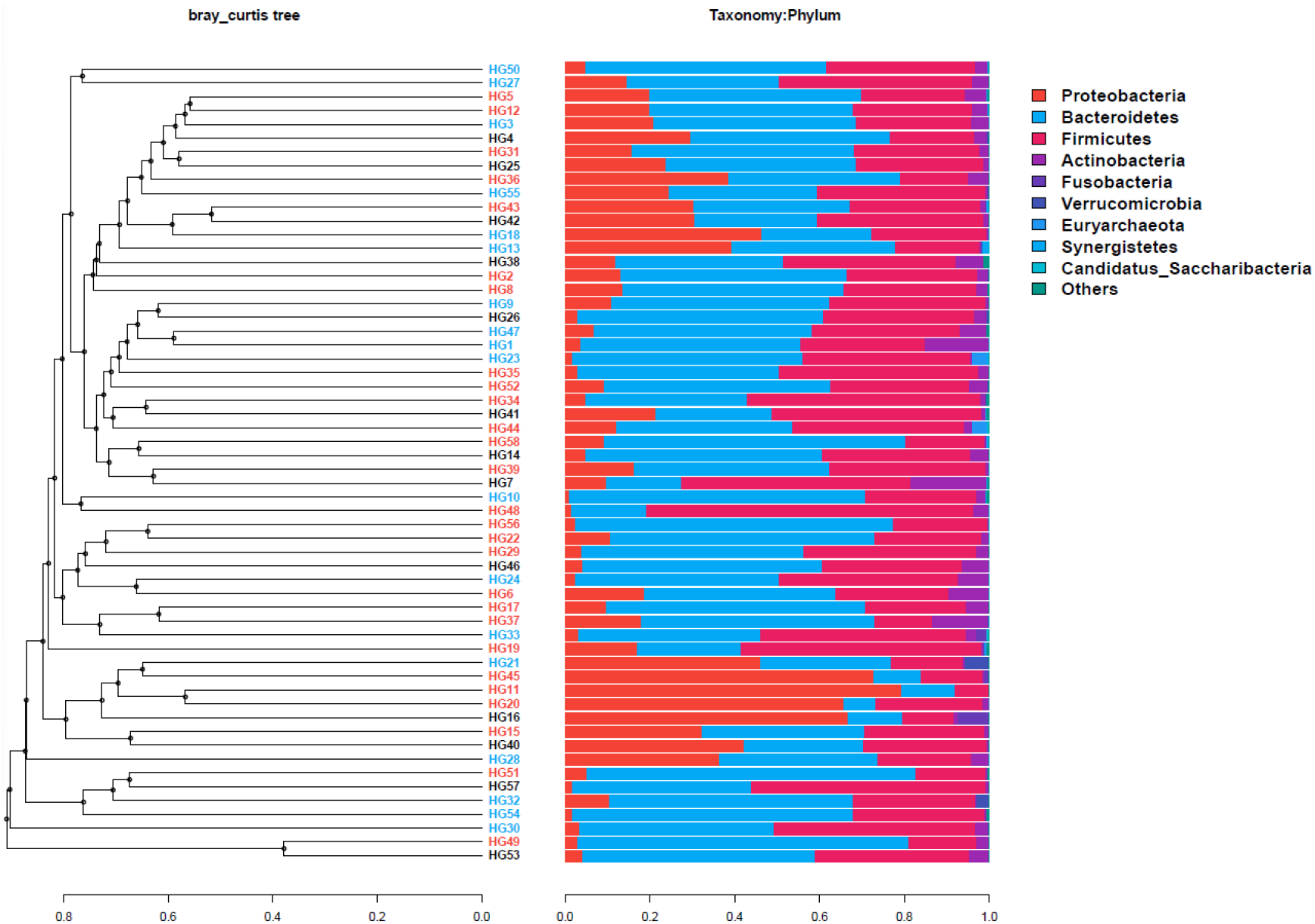
Beta diversity distribution of gut microbe samples from Zhejiang and Shanghai

In the three groups, the proportion of *Bifidobacterium* was 2.72% (TH group), 2.20% (MI group) and 2.03% (FA group), respectively. *Bifidobacterium* is a recognized genus of probiotics, which can help digestion and decomposition, regulate immune response and inhibit pathogens [15]. It decreased with the increase of BMI, indicating that *Bifidobacterium* may help maintain human health. In overweight people, the number of *Bifidobacteria* may decrease due to metabolic reasons. The abundance of *Lactobacillus* in underweight group and overweight group was 0.11% and 0.16%, respectively, which was lower than that in the normal group by 0.29%. *Lactobacillus* is a common intestinal probiotic that converts sugars into lactic acid through fermentation. It can reduce the risk of coronary heart disease, alleviate the symptoms of irritable bowel syndrome, and regulate the balance of intestinal microbes. Studies have shown that moderate intake of lactic acid bacteria in healthy people can increase the content of short-chain fatty acids in intestinal tract and maintain intestinal health [20].

*Helicobacter* was also detected in 19 samples, suggesting that these populations may be infected with *Helicobacter pylori* [11] [25]. In addition, some samples also contain *Klebsiella*, a genus of microbes often associated with drug-resistant pathogens, therefore it was also suggested that potential infections from those bacteria should be concerned. In addition to these dominant and potentially pathogenic bacteria genera, there are hundreds of other genera in these gut microbiota. Ongoing efforts are still needed to gain a better understanding of the human gut microbiota.

### 3.5. Analysis of dominant bacteria based on OTU

At the OTU level, the top 20 abundant ZOTU species accounted for 34.11% of all sequences, indicating the existence of dominant species in these communities. The average distribution of *Escherichia* OTU in the gut microbiota of 58 individuals was 6.11%, and the proportions in TH, MI and FA groups were 7.74%, 6.38% and 4.53%, respectively, indicating that the OTU decreased with the increase of BMI (Table 2). Among these populations, *Bacteroidetes* had more dominant OTU, such as ZOTU_2 and ZOTU_3. Previous studies have shown that ZOTU_2 predicted *Bacteroides Uniformis* was related to inflammation. In obesity caused by high-fat diet, this bacteria can improve metabolic and immune disorders and maintain human health [22]. The distribution of the bacteria was the lowest in the TH group, only 3.06%, but its proportion in the MI and FA groups was 4.57% and 4.06%, respectively, further suggesting that the bacteria may be associated with BMI (Table 2). The abundance of *Bifidobacterium* was distributed differently among the three groups. At the OTU level, the most dominant OTU of *Bifidobacterium* was OTU24, which has an average proportion of 0.68% in all samples. Its distribution in the three groups was 0.62%, 0.54% and 0.95%, respectively (Table 2). The proportion of this strain in overweight population is relatively higher, indicating that it is possible to screen microbial functional species related to obesity and other diseases by analyzing the strain at the level of subdivided OTU. *Akkermansia muciniphila* is a mucin-breaking bacterium that has been shown to be associated with human health, particularly in athletes. However, the abundance of ZOTU_142 was 0.33% in the overweight FA group, but was lower in the TH and MI groups, and even not detected. In particular, the abundance of *Akkermansia* in two samples from the overweight group was high, reaching 5.62% and 2.86%. Further research on these two samples is expected to better explain the correlation between the distribution characteristics of microbes in different people and BMI and human health status. *Akkermansia muciniphila* is a recently discovered probiotics, which has been shown to be abundant in the milk of lean and non-diabetic women, and is speculated to be a potential solution for metabolic diseases (such as type II diabetes and obesity) [21].

**Table 2.** OTU analysis of gut microbiota dominance in Zhejiang and Shanghai

|  | TH | MI | FA | Predicted taxonomy |
| --- | --- | --- | --- | --- |
| ZOTU_1 | 7.74% | 6.38% | 4.53% | g:Escherichia/Shigella |
| ZOTU_2 | 3.06% | 4.57% | 4.06% | Bacteroides uniformis |
| ZOTU_3 | 1.16% | 3.28% | 1.44% | Bacteroides dorei |
| ZOTU_7 | 2.12% | 1.34% | 2.41% | g:Bacteroides |
| ZOTU_8 | 3.12% | 1.40% | 1.11% | Bacteroides uniformis |
| ZOTU_5 | 1.04% | 1.59% | 2.05% | g:Alistipes |
| ZOTU_6 | 1.73% | 1.24% | 1.92% | Faecalibacterium prausnitzii |
| ZOTU_4 | 2.50% | 0.77% | 1.33% | Clostridiales |
| ZOTU_9 | 0.67% | 2.03% | 0.63% | f:Enterobacteriaceae |
| ZOTU_11 | 1.15% | 0.91% | 1.94% | Blautia wexlerae |
| ZOTU_10 | 0.65% | 1.58% | 1.13% | f:Lachnospiraceae |
| ZOTU_14 | 0.93% | 1.17% | 1.54% | g:Bacteroides |
| ZOTU_13 | 0.97% | 1.53% | 0.93% | Phascolarctobacterium faecium |
| ZOTU_25 | 1.47% | 0.56% | 2.00% | Bacteroides plebeius |
| ZOTU_12 | 0.49% | 0.96% | 1.67% | Parabacteroides merdae |
| ZOTU_22 | 0.37% | 1.10% | 1.53% | g:Bacteroides |
| ZOTU_20 | 1.81% | 1.05% | 0.46% | g:Bacteroides |
| ZOTU_19 | 0.21% | 1.28% | 1.21% | f:Enterobacteriaceae |

### 3.6. Beta diversity analysis of gut microbiome samples

The beta diversity of the community can reflect the relationship between samples. After classifying the relevant samples applying various beta analysis methods, it was found that the differences among the samples in the three groups were not significant, resulting in the samples within different groups being close to each other. However, it can be seen from the results that some samples from the same group are relatively close, such as HG6, HG17 and HG37. This indicates that the grouping of all samples by BMI should be improved due to other influencing factors. In addition, people of different ages in the same family were involved in this study, and the difference in gut microbiota was interfered by many different factors, such as diet and age, making it difficult to analyze the influencing factors of BMI alone. In future studies, the gut microbiota of individuals with large BMI differences in the same age, gender and region should be strictly selected, the possible association between BMI value and gut microbiota should be analyzed separately to reduce the influence of other factors on the relationship between BMI value and gut microbiota.

### 3.7. Isolation and culture of some microorganisms in gut microbiota

LB and MRS medium were used to isolate and culture gut bacteria from a healthy teenager, and a total of 17 monoclones were obtained. Some microbial DNA from these clones were extracted according to the colonial morphology and mycelial morphology, and 4 of them were sequenced. After comparing blastN with the nucleic acid sequence database in NCBI database, the four microbial strains were found, one was *Escherichia fergusonii* and the other three were *Weissella cibaria* (Table 3).

**Table 3.** Comparison results of isolated microorganisms with those in NCBI database

|  | Closest isolated strain (serial number) | Similarity index |
| --- | --- | --- |
| HX-111 | *Escherichia fergusonii* strain ATCC 29903 (NR_026331.1) | 99.49% |
| HX-251 | *Weissella cibaria* strain II-I-59 (NR_036924.1) | 99.79% |
| HX-261 | *Weissella cibaria* strain II-I-59 (NR_036924.1) | 99.71% |
| HX-263 | *Weissella cibaria* strain II-I-59 (NR_036924.1) | 99.79% |

*Escherichia fergusonii* exists mostly in the environment and animal intestines which is an opportunistic pathogen as well as a common bacterium in the intestinal tract. Escherichia was also the second most abundant microbial group of this research in sequencing, indicating that a dominant strain of intestinal bacteria was also obtained through isolation and culture (Figure 5 and Figure 6). *Weissella cibaria* is a lactic acid bacterium that colonized in the gut, belonging to the phylum Firmicutes and is a probiotic bacterium. The bacteria protect the host by producing organic acids to resist bacteria and prevent pathogens from attaching to the intestinal epithelium. At the same time, the bacteria can scavenge free radicals in the intestinal tract and inhibit lipid peroxidation, exhibiting antioxidant effects [13]. However, the abundance of this bacterium was low in high-throughput sequencing and only existed in part of human gut microbiota, indicating that the non-dominant probiotics were isolated using probiotic medium MRS. In future research, according to the results of next-generation sequencing, a suitable medium can be selected or designed, and the dominant microbial groups in it can be isolated and cultured to provide strains for studying the relationship between bacterial groups and BMI values and other potential diseases. In addition, this can provide population-derived strains in the region for microbiota regulation.

## 4. Conclusion

After analyzing the gut microbiota samples of people with different BMI values in Zhejiang and Shanghai, the results have shown that the population in this area was different from the known enterotypes, which had a high degree of novelty. Therefore, it was necessary to try to establish the gut microbiota database of different regions. Also, the Firmicutes/Bacteroidetes ratio increases with the decrease of BMI, which conforms to previous results [8]. The analysis of these populations showed that the gut microbiota contained high *Bacteroides*, indicating that the population in this area mainly consumed protein food. To maintain health, relevant diets should be adjusted to increase the proportion of Firmicutes. In addition, four strains were isolated from intestinal bacteria, belonging to high-abundance *Escherichia* and low-abundance *Weisseria cibaria*, respectively, indicating that isolation and culture of microorganisms is expected to provide a material basis for the regulation of gut microbiota in the future.

It’s necessary to clarify that the sample size of this study was small, with large differences in age among people from Zhejiang and Shanghai. Although it was also found that the reduction of *Bifidobacteria* and other factors may be related to high BMI, there was no significant difference among people with different BMI values. In future studies, it is necessary to further expand the sample size and select the population of the same region, age, and gender to reduce the potential impact of age, regional dietary differences and other factors on gut microbiota.

## Acknowledgement

This paper was written under the guidance of Ms. Lv Yinghe. Here, I would like to apprecia her work and careful guidance. At the same time, I would like to thank Associate Professor Wei Yongjun of Zhengzhou University for his technical support and guidance.

